# Highly accurate long reads are crucial for realizing the potential of biodiversity genomics

**DOI:** 10.1101/2022.07.10.499467

**Authors:** Scott Hotaling, Edward R. Wilcox, Jacqueline Heckenhauer, Russell J. Stewart, Paul B. Frandsen

**Affiliations:** Department of Watershed Sciences, Utah State University, Logan, UT, USA; DNA Sequencing Center, Department of Biology, Brigham Young University, Provo, UT, USA; LOEWE Centre for Translational Biodiversity Genomics (LOEWE-TBG), Frankfurt, Germany; Department of Terrestrial Zoology, Senckenberg Research Institute and Natural History Museum Frankfurt, Frankfurt 60325, Germany; Department of Biomedical Engineering, University of Utah, Salt Lake City, UT, USA; Department of Plant and Wildlife Sciences, Brigham Young University, Provo, UT, USA; Data Science Lab, Smithsonian Institution, Washington, DC, USA

**Keywords:** Insecta, Oxford Nanopore, PacBio, HiFi, caddisfly, genome biology

## Abstract

**Background:** Generating the most contiguous, accurate genome assemblies given available sequencing technologies is a long-standing challenge in genome science. With the rise of long-read sequencing, assembly challenges have shifted from merely increasing contiguity to correctly assembling complex, repetitive regions of interest, ideally in a phased manner. At present, researchers largely choose between two types of long read data: longer, but less accurate sequences, or highly accurate, but shorter reads (i.e., >Q20 or 99% accurate). To better understand how these types of long-read data as well as scale of data (i.e., mean length and sequencing depth) influence genome assembly outcomes, we compared genome assemblies for a caddisfly, *Hesperophylax magnus*, generated with longer, but less accurate, Oxford Nanopore (ONT) R9.4.1 and highly accurate PacBio HiFi (HiFi) data. Next, we expanded this comparison to consider the influence of highly accurate long-read sequence data on genome assemblies across 6,750 plant and animal genomes. For this broader comparison, we used HiFi data as a surrogate for highly accurate long-reads broadly as we could identify when they were used from GenBank metadata.

**Results:** HiFi reads outperformed ONT reads in all assembly metrics tested for the caddisfly data set and allowed for accurate assembly of the repetitive ∼20 Kb *H-fibroin* gene. Across plants and animals, genome assemblies that incorporated HiFi reads were also more contiguous. For plants, the average HiFi assembly was 501% more contiguous (mean contig N50 = 20.5 Mb) than those generated with any other long-read data (mean contig N50 = 4.1 Mb). For animals, HiFi assemblies were 226% more contiguous (mean contig N50 = 20.9 Mb) versus other long-read assemblies (mean contig N50 = 9.3 Mb). In plants, we also found limited evidence that HiFi may offer a unique solution for overcoming genomic complexity that scales with assembly size.

**Conclusions:** Highly accurate long-reads generated with HiFi or analogous technologies represent a key tool for maximizing genome assembly quality for a wide swath of plants and animals. This finding is particularly important when resources only allow for one type of sequencing data to be generated. Ultimately, to realize the promise of biodiversity genomics, we call for greater uptake of highly accurate long-reads in future studies.

## Introduction

As genome sequencing has been revolutionized by high-throughput sequencing, a general rule has emerged: more data yields more contiguous, accurate genome assemblies. This is particularly evident when read length is considered; third-generation long reads, which are often tens or even hundreds of thousands of base pairs in length, have dramatically improved genome assemblies across the Tree of Life (1-4). For coverage, an increase can limit the impacts of erroneous read calls through more replication of potential variants (5). However, a shortcoming of second-generation, short-read platforms (e.g., Illumina) for genome assembly is that no amount of data will allow for resolution of repeat-driven gaps that exceed read lengths (5). The power of long reads to mitigate this issue is well-documented (e.g., 6, 7, 8) but not all long reads are created equal; different platforms, technologies, and chemistries yield different read length versus error profiles (9-11). This difference is particularly important since some long, repeat-rich genomic regions–including many genes of phenotypic relevance (e.g., antifreeze proteins in polar fishes)–pose assembly challenges even when long read data are used (6).

Generally speaking, past evidence suggests that to resolve difficult genomic regions with long read data, longer reads at greater depth of coverage will outperform shorter reads and/or less dense coverage (12). However, it is unclear how this expectation jibes with read accuracy, particularly for two common types of long-read data that are currently in use: longer but noisier long reads and more accurate, but shorter, long reads.

We tested this prediction—that longer, higher coverage, and noisier reads would outperform shorter, but more accurate, reads at lower sequencing depth—in a silk-producing caddisfly, *Hesperophylax magnus*, with an emphasis on the highly repetitive *heavy fibroin chain* gene *(H-fibroin*). Caddisflies share an evolutionary origin of silk with butterflies and moths, including the primary protein component of silk, *H-fibroin* (13). *H-fibroin* commonly spans ∼20 Kb and its amino acid sequence consists of conserved termini and a highly repetitive internal region. Early efforts to assemble the complete *H-fibroin* gene with short reads were unsuccessful due to its long repetitive region (14, 15). However, long-read assemblies have since yielded full-length sequences (13, 16, 17).

We compared two long-read genome assemblies for *H. magnus* produced with two technologies: earlier generation Oxford Nanopore (ONT) R9.4.1 reads and more accurate PacBio HiFi (HiFi) reads from the same era. For both assemblies, we considered genome-wide metrics as well as accurate assembly of the *H-fibroin* gene. We focused on *H-fibroin* as a surrogate for complex but phenotypically important genes where we expected the benefits of highly accurate long reads to be most obvious. To estimate whether assembly outperformance with highly accurate reads (i.e., >Q20 or 99% accurate) was unique to our focal caddisfly or reflects a broader trend in genome biology, we performed a meta-analysis of contig N50, assembly length, and sequencing technology for all publicly available plant and animal genomes on GenBank. For this analysis, it should be noted that while we focused on HiFi data due to methodological limitations, recent ONT technologies have also reached this accuracy threshold. For all of our evaluation metrics (e.g., contiguity, BUSCO, accurate assembly of complex regions), highly accurate long-read sequence data appears to dramatically outperform other types of sequence data.

## Results

For our caddisfly genome assembly comparison, the ONT library had a wider distribution of read lengths with a median of 19.9 Kb for 33.5 Gb of raw data (27.2x coverage). The HiFi dataset had a median read length of 11.3 Kb for 28 Gb of raw data (22.8x coverage; Fig. 1). The best ONT assembly (Genbank #GCA_016648045.1) was generated with MaSuRCA and spanned 1.23 Gb. For HiFi, the best assembly was produced with Hifiasm (Genbank #JAIUSX000000000) and was nearly identical in length at 1.22 Gb. However, we observed a dramatic difference in contiguity; the ONT assembly had a contig N50 of 0.7 Mb versus 11.2 Mb for the HiFi assembly. The ONT assembly also contained fewer complete BUSCOs (93%) versus the HiFi assembly (95.6%). In both assemblies, the full-length *H-fibroin* locus was present but the quality of the annotations differed greatly (Fig. 2). For ONT, we annotated a dozen genes in the ∼30 Kb region, most of which did not include the characteristic repeats known from previous data. For HiFi, the annotation included a single gene with a single intron in the n-terminus region. The second exon was large (25.8 Kb) and fully in-frame including a well-resolved repetitive structure, giving high confidence in the accuracy of the assembly.

**Figure 1.**
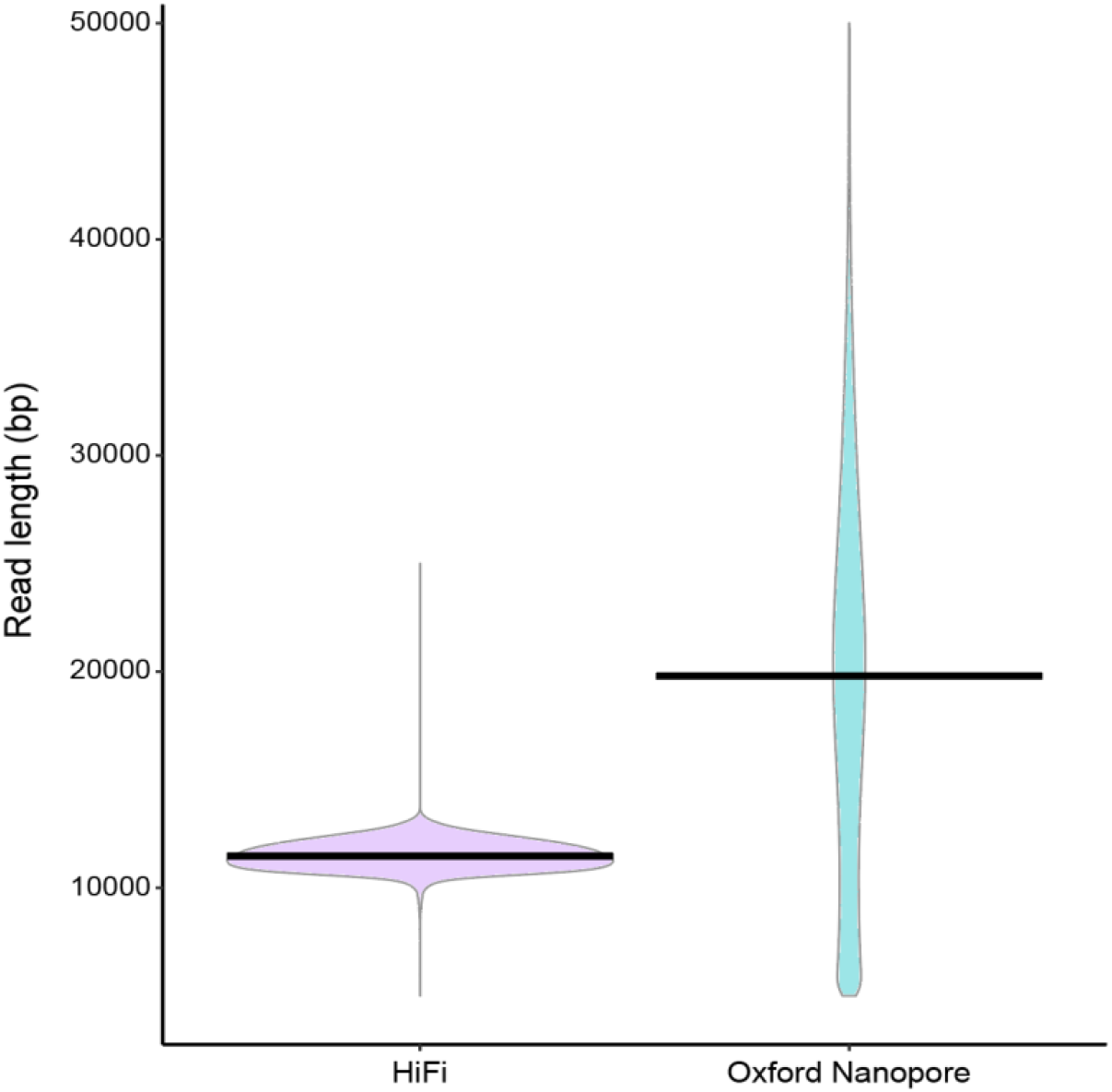
Violin plots of read lengths for the HiFi and Oxford Nanopore data sets used to assemble *Hesperophylax magnus* genomes in this study. Width of the colored areas indicate numbers of reads according to lengths on the y-axis. Dark lines represent the medians of each distribution.

**Figure 2.**
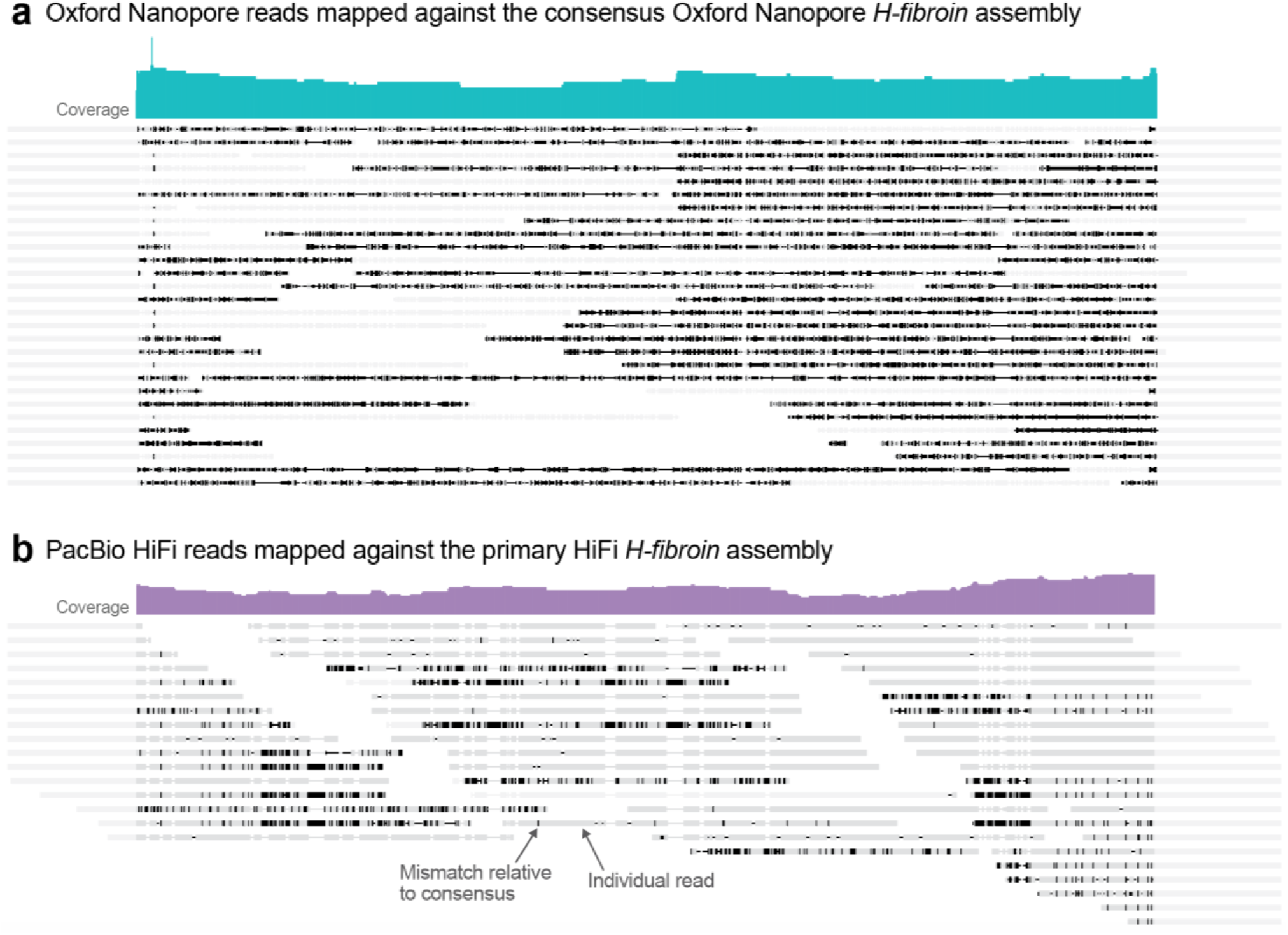
A case study comparing the capacity for two long-read sequencing technologies to assemble the complex gene underlying silk production in caddisflies, *H-fibroin*. (a) Raw Oxford Nanopore (ONT) reads mapped to the consensus *H-fibroin* sequence from the ONT assembly. (b) Raw HiFi reads mapped to the primary *H-fibroin* sequence from the phased HiFi assembly. Dark lines indicate mismatches relative to consensus. In (b), mismatches reflect an *H-fibroin* length polymorphism that can be resolved by subsampling reads based on their allele-specificity. Note the high amount of noise within reads relative to the consensus for highly accurate long reads (b) versus longer, but less accurate, reads (a).

After filtering, our meta-analysis data set contained 6,750 genome assemblies (animals = 5,592; plants = 1,158; Table S1). For plants, short-read assemblies (48.1%; *N* = 557) and long-read assemblies generated with non-HiFi PacBio data were similarly common (39.6%; *N* = 458; Fig. 3a). ONT assemblies, however, were much less common (10.7%) and HiFi assemblies were exceptionally rare, comprising just 1.6% of all assemblies (*N* = 19; Fig. 3a). For animals, the majority of assemblies were generated with short-read data (74.1%; *N* = 4,142). Non-HiFi long-read assemblies, generated with either PacBio or ONT reads, were also common, comprising 21.4% of the data set. HiFi assemblies were again the least common with just 259 assemblies (4.6%; Fig. 3b).

**Figure 3.**
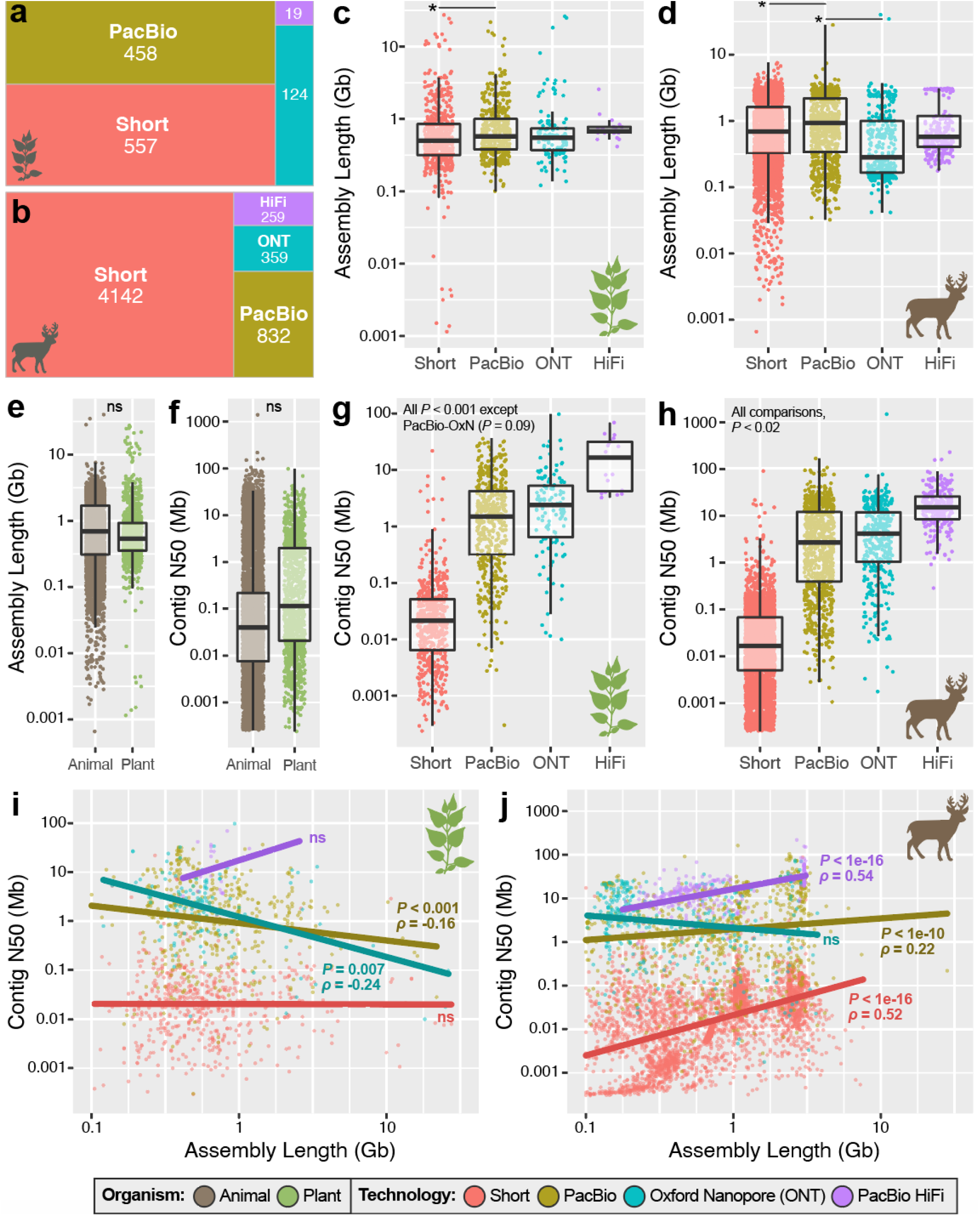
Sequencing technology representation and genome assembly quality across all animal and plant assemblies deposited in GenBank as of November 2021. A breakdown of the sequencing technology used for genome assemblies in (a) plants and (b) animals. Total assembly length broken down by sequencing technology for (c) plants and (d) animals. (e) Assembly length across all plant and animal genomes, regardless of technology. (f) Contig N50 across all plant and animal genomes and broken down by technology for (g) plants and (h) animals. Spearman’s correlations between contig N50 and assembly length for (i) plants and (j) animals. For (i) and (j), correlation statistics were generated for the full data sets but for visualization, assemblies less than 0.1 Gb in length or with contig N50 > 1 Gb have been excluded. For (c-h), asterisks and thin dark lines indicate significant differences at *P* < 0.05.

On average, animal genome assemblies were neither longer (*P*, Welch T-test = 0.80, Fig. 3e) nor more contiguous than those of plants (*P*, Welch T-test = 0.10, Fig. 3f). When broken down by technology, assembly lengths did differ for plants (*P*, one-way ANOVA = 0.006) and animals (*P*, one-way ANOVA < 0.001). For plants, only one length comparison was significantly different–non-HiFi PacBio assemblies were longer than those generated with short-reads (*P*, Tukey HSD = 0.005; Fig. 3c). For animals, non-HiFi PacBio assemblies were longer than both short-read (*P*, Tukey HSD = 0.002) and ONT assemblies (*P*, Tukey HSD < 0.001; Fig. 3d).

In terms of assembly contiguity, contig N50 was significantly different for all comparisons in plants (*P*, Tukey HSD < 0.001) and animals (*P*, Tukey HSD < 0.02) with one exception: in plants, contiguity of non-HiFi PacBio assemblies was not different from ONT assemblies (*P*, Tukey HSD = 0.09; Fig. 3g-h). For both groups, highly accurate HiFi reads dramatically outperformed all other long-read technologies tested. In plants, the average HiFi assembly was 501% more contiguous (mean contig N50 = 20.5 Mb) than assemblies generated with other long-reads (mean contig N50 = 4.1 Mb; Fig. 3g). For animals, HiFi assemblies were 226% more contiguous (mean contig N50 = 20.9 Mb) versus other long-read assemblies (mean contig N50 = 9.3 Mb; Fig. 3h). To assess if recent ONT technological improvements (e.g., R10.4) yielded more contiguous assemblies than our broader analysis captured, we performed an additional analysis focused on only assemblies from 2021. We found no difference between ONT and non-HiFi PacBio assembly contiguity for this most recent subset of the animal data (*P*, one-way ANOVA = 0.26).

When assembly size was compared to contiguity, clear patterns emerged. For plants, when non-HiFi long reads are used, contiguity declines with increasing assembly length (*P* < 0.008; Spearman’s ⍴, PacBio = -0.16, Spearman’s ⍴, ONT = -0.24; Fig. 3i). The same trend isn’t present for short reads (*P*, Spearman’s ⍴ = 0.18) nor HiFi (*P*, Spearman’s ⍴ = 0.37). However, for HiFi, this lack of significance is likely a product of small sample size (*N* = 19; Fig. 3i). For animals, however, three of four read types (short, PacBio, HiFi) exhibit positive correlations between contig N50 and assembly length (*P*, Spearman’s ⍴ < 1e-10) with the steepest trends for the two most accurate sequencing technologies: HiFi (Spearman’s ⍴ = 0.54) and short-reads (Spearman’s ⍴ = 0.52; Fig. 3j).

## Discussion

As high-throughput sequencing technologies have matured, a common strategy has emerged: to generate better assemblies, more data is better. While simplistic, this “more is better” approach has been supported by empirical data and echoed by genome sequencing overviews (e.g., 18). The practicality of this approach has also been empirically observed in the long-read era. For instance, benchmarking of long-read assemblies in maize found that lower sequencing depths (< 30x) with mean read lengths less than 11 Kb yielded highly fragmented assemblies (12). Stepping back, the outperformance of long-reads relative to short-reads in terms of basic assembly metrics is dramatic and independent of taxonomy (1, 3). Thus, support exists for the premise that to generate the most high-quality assemblies, researchers should maximize depth of coverage and mean read length regardless of the technology being used.

However, since we cannot maximize perfectly accurate deep sequencing with the longest possible reads, particularly when limited resources preclude hybrid approaches, achieving the best possible genome assemblies under the current landscape of sequencing technologies may require more nuance. Our results highlight that in at least one species, a smaller amount of shorter, but more accurate, HiFi reads outperformed ONT data for genome assembly. Given the scale of outperformance—approximately an order of magnitude when contiguity is considered— it is clear that biological variation between individuals from the same population would not account for this result. Rather, this outperformance likely stems from base pair accuracy and the potential for sequence errors to confound assembly. Thus, we expected the benefit of highly accurate reads to scale with genome and/or gene-region complexity. The dramatic outperformance of highly accurate HiFi reads when assembling the *H-fibroin* locus supports this expectation. Similar results have been obtained for other taxonomic groups. For the leaf rust fungus, HiFi reads outperformed noisier long-reads, particularly in areas of the genome that were difficult to assemble (19). Among rice genomes, however, the results were less clear: ultra-long ONT reads yielded higher overall contiguity but more errors than HiFi (20). Notably, when resources allow for two long-read data sets to be generated, hybrid assemblies that integrate, for instance, ultra-long reads such as those generated with PacBio continuous long read (CLR) technology combined with highly accurate long reads (e.g., HiFi) can improve results (21, 22).

Moving beyond our case study to a large-scale comparison of animal and plant genome assemblies, additional themes emerged. First, assemblies generated with highly accurate long-read technologies—in this case, HiFi—were significantly more contiguous than all other types of sequence data tested. Despite this, HiFi sequencing remains underrepresented, particularly in plant genetics. Second, all significant relationships between genome size and contiguity for sequencing technologies in animals were positive (Fig. 3j) whereas no correlation was positive in plants (Fig. 3i). This suggests that as genome size increases in plants, so too does complexity, likely at a rate that outpaces the capacity for modern assembly algorithms to assemble it. Notable, however, was the sharply positive trend for HiFi sequencing in plants where contig N50 appears to rapidly increase with assembly length (Fig. 3i). While not statistically significant at *P* < 0.05–likely due to a low sample size–this pattern suggests that HiFi and similarly accurate long-read technologies may represent a valuable means to overcome challenges of genome complexity in plants.

However, despite the striking patterns we observed, we must acknowledge that our approach inherently biases our conclusions towards HiFi as this was the only highly accurate long-read technology that could be distinguished from the available metadata. Indeed, GenBank assemblies commonly reported when HiFi reads were incorporated or when a HiFi-specific assembler was used. No such reporting exists for the type of ONT data (e.g., R9 vs. R10) nor are there highly accurate ONT-specific assemblers that can be used as surrogates for this information. Thus, we want to be clear that while we focused on specific technologies in this study and found strong evidence for HiFi efficacy, our point is more about long-read accuracy broadly than specific technologies. Indeed, other technologies–including ONT–have approached and may even surpass HiFi in terms of accuracy in the months and years to come. Finally, to close this data gap for future meta-analyses, we echo previous studies that have called for improvements to the metadata reporting process on GenBank (2).

With the rise of highly accurate long-read sequencing, little room remains for another revolution in genome assembly quality. Relative to even a few years ago, it is now possible to generate reference-quality assemblies for virtually any species with modest resources (e.g., less than $5,000 for a 1-2 Gb assembly). The only exceptions, for now, are species that are very small, difficult to obtain, or with exceptionally large genomes. Now, we lack appropriate metrics for contemporary genome assembly benchmarking. For instance, contig N50–the most common metric for assessing contiguity–scales with assembly length in highly accurate long-read assemblies (Fig. 3i-j) and thus, its upper limit is tied to chromosome length, making comparisons among groups difficult. For gene content assessment–i.e., BUSCO scores–one challenge lies in how accurately BUSCO scores reflect true gene content, particularly when more repeat-rich genes are considered. For instance, the median gene length in the 1,467-gene “Insecta” gene set is ∼1 Kb (longest = 9.1 Kb; 1) yet phenotypically relevant genes like *H-fibroin* can be *much* longer (i.e., >20 Kb). Thus, while BUSCO scores for the ONT and HiFi assemblies of *H. magnus* reflect marginal differences in their gene completeness (93% vs. 95.6%), the true gap is likely much greater. Indeed, we expect differences in assembly quality to scale with genomic region complexity–a result that is not captured by BUSCO scores.

## Methods

For our caddisfly genome comparison, we sequenced two individuals of *H. magnus* from the same population. High-molecular weight DNA was extracted using the same method for both individuals (Agilent DNA Extraction kits). For the first individual, we generated a combination of noisy ONT sequencing (MinION R.9.4.1 flow cells with the LSK-109 ligation library prep kit) and Illumina sequencing (PE 150 bp, ∼49x coverage; Illumina NovaSeq 6000). Based on published ONT benchmarking (https://nanoporetech.com/accuracy), we expect our ONT reads had a raw read accuracy of ∼98.3%. For an extended description of the ONT sequencing and assembly methods, see the associated publication (23). For the second individual, we only generated HiFi reads via circular consensus sequencing (CCS) on the PacBio Sequel II platform. While use of the same individual for both libraries would have been an ideal way to control for biological noise, the reality of extracting large amounts of high molecular weight DNA for long-read sequencing made this difficult for our study. For PacBio technologies, we refer to HiFi reads as those generated with CCS and “non-HiFi” reads as anything else generated on a PacBio platform. By definition, CCS HiFi reads have raw read accuracy >Q20 (>99%).

We used Guppy v.5.0.11 in the “high accuracy” mode to base-call the ONT reads and removed all reads under 5 Kb for further analysis. We also filtered reads for quality using default settings in MinKNOW. Next, we used SMRTlink v.10 to generate HiFi reads (reads with quality >Q20). To assemble genomes for the two individuals, we tested a range of assemblers. For ONT, we tested Flye v.2.9.1 (24), Canu v.1.8 (25), wtdbg2 v.2.4 (26), and a hybrid approach with MaSuRCA that integrated both ONT and Illumina short-read data (27). For HiFi, we tested Flye v.2.9.1 (24), Hifiasm (28) and HiCanu (29). We selected the best assembly based on contiguity (contig N50) and “Benchmarking Universal Single-Copy Orthologs” (BUSCO) scores. We ran BUSCO v.5.2.2 (30) using the 1,367 reference genes in the OrthoDB v.10 Insecta gene set (31). We phased the best HiFi assembly, which was generated with Hifiasm (see Results), using the program’s built-in phasing which yielded primary and alternate assemblies. No auto-phasing method is present for MaSuRCA (which produced the best ONT assembly). No post-assembly polishing of the MaSuRCA assembly was performed (23).

To evaluate recovery and assembly of the *H-fibroin* locus, we used tblastn to identify conserved terminal sequences from existing transcriptomes (14). We then extracted *H-fibroin* and 1,000 additional bps from each terminus from both assemblies and annotated the region using Augustus v.3.3.2 (32). For comparison to the HiFi assembly, we attempted to phase the best ONT assembly using WhatsHap (33) for the *H-fibroin* locus but failed due to the high number of mismatches with the consensus sequence which meant that only a few reads could be assigned to a haplotype. To visualize mismatches between reads and both assemblies, we mapped raw reads to the assembled *H-fibroin* gene using Minimap2 (34) and visualized the results in Geneious 2022.0.2 (35).

To assess the influence of sequencing technology on genome assembly across the Tree of Life, we extracted metadata for all plant (class Embryophyta) and animal (class Metazoa) genome assemblies from GenBank using the “summary genome” function in v.10.9.0 of the NCBI Datasets command-line tool on 13 November 2021. We then used the “lineage” function of TaxonKit (36) to retrieve taxonomic information for all entries in our genome assembly list. Next, we gathered additional metadata (e.g., sequencing technology) for each entry using a custom web scraper script (modified from 1). We removed duplicate assemblies and alternative haplotypes for a given assembly through keyword searching in either the BioProject information or assembly title. For this meta-analysis, we used contig N50 as a correlate for genome assembly quality. We acknowledge that while this is a useful, widely used metric for assessing genome assembly quality [e.g., (37)], it is not without flaws including the reality that it scales with chromosome length and is therefore difficult to compare across taxa.

We binned assemblies into four sequencing technology categories: short-reads (e.g., Illumina), long-read ONT (ONT long-reads with or without short-reads), long-read PacBio (non-HiFi PacBio long-reads with or without short reads), or HiFi (any assembly where HiFi long-reads were used). To assist in categorization, we considered any assembly that was generated before 2017—when long-read assemblies began to emerge (1)—to be a short-read assembly. We also used self-reported information on the genome assembly algorithms used to classify assemblies. For instance, if an assembly only reported PacBio sequence data but a HiFi-specific assembler (e.g., Hifiasm) was used, we classified it as a HiFi assembly. For this comparison, we emphasized HiFi reads as our surrogate for highly accurate long reads because it was impossible to distinguish types of ONT reads (e.g., different pores, chemistries, etc.) from the GenBank metadata. We removed any assembly for which the sequence data type used could not be established.

We tested for differences in the distributions of assembly size and contig N50 among plant and animals or our sequence type categories within the overarching plant or animal grouping using Welch Two Sample T-tests or one-way ANOVAs followed by Tukey HSD tests in R v3.6.3 (38). While all statistical tests were performed on the untransformed data, we visualized log-transformed comparisons using ggplot2 (39). Because we were only able to distinguish one type of highly accurate long read data (i.e., HiFi) from other long-read data types, we performed an additional comparison between ONT and PacBio (non-HiFi) assemblies for the 2021 animal data. The goal of this analysis was to assess if a recent shift in ONT assembly quality was occurring in parallel with new technological developments (specifically R10.4) that we could be overlooking with our general binning across years.

While our high-level approach for assessing the impacts of HiFi on genome assembly metrics for plants and animals broadly may be confounded by other factors (e.g., genome size variation across groups), the small number of HiFi assemblies within any particular group—particularly at the level where genome size variation, for instance, is minimized (e.g., genus)—limited our capacity to make more focused comparisons. Further, even if robust within groups comparisons could be made (e.g., within humans), it is unclear how well these comparisons would reflect patterns in non-model groups without a history of high-quality reference sequences for assembly.

## Declarations

### Ethics approval and consent to participate

Not applicable.

### Consent for publication

Not applicable.

### Availability of data and materials

The dataset generated and analyzed for the animal and plant genome analysis is available in Table S1. Additionally, the genome assemblies and raw data are available in the Genbank repository (https://www.ncbi.nlm.nih.gov/genbank/) with accession numbers (#GCA_016648045.1, #JAIUSX000000000) and the raw sequence data accession numbers are: ONT = SRR12822505, Illumina = SRX9290147, HiFi = SRR15840267.

### Authors’ contributions

S.H and P.B.F designed the study, analyzed data, and wrote the manuscript with input from all coauthors. E.R.W. performed laboratory work and generate the sequence data for both caddisfly assemblies used. J.H. and R.J.S. contributed to caddisfly data generation and provided input on manuscript development and content.

## Acknowledgements

J.H was supported by the LOEWE-Centre for Translational Biodiversity Genomics, which was funded by the Hessen State Ministry of Higher Education, Research and the Arts.

